# High-content cellular screen image analysis benchmark study

**DOI:** 10.1101/2022.05.15.491989

**Authors:** Mark-Anthony Bray, Antoine de Weck, Eric Durand, Jian Fang, Daniela Gabriel, Rens Janssens, Ioannis Moutsatsos, Stephan Spiegel, Xian Zhang

**Affiliations:** Novartis Institutes for BioMedical Research Inc., Cambridge, MA, USA; Novartis Institutes for BioMedical Research, Basel, Switzerland; Children’s Cancer Institute, Lowy Cancer Research Centre and School of Women and Children’s Health, Faculty of Medicine and Health, UNSW Sydney, NSW, 2031, Australia

## Abstract

Recent development of novel methods based on deep neural networks has transformed how high-content microscopy cellular images are analyzed. Nonetheless, it is still a challenge to identify cellular phenotypic changes caused by chemical or genetic treatments and to elucidate the relationships among treatments in an unsupervised manner, due to the large data volume, high phenotypic complexity and the presence of a priori unknown phenotypes. Here we benchmarked five deep neural network methods and two feature engineering methods on a well-characterized public data set. In contrast to previous benchmarking efforts, the manual annotations were not provided to the methods, but rather used as evaluation criteria afterwards. The seven methods individually performed feature extraction or representation learning from cellular images, and were consistently evaluated for downstream phenotype prediction and clustering tasks. We identified the strengths of individual methods across evaluation metrics, and further examined the biological concepts of features automatically learned by deep neural networks.

## Introduction

High-content cellular imaging assays stain cells with multiple dyes and fluorescent-labelled antibodies, visualize cellular compartments and organelles as well as specific proteins in high-resolution multiple-channel microscopy images, and provide rich cellular phenotypic information. In a high-throughput mode, high-content cellular imaging assays can characterize phenotypic changes caused by individual genetic or chemical perturbations of a large library, and thus are a powerful technology widely applied to investigation of the functional effects of genes and compounds for biological research and drug discovery^1–3^. The imaging data acquired from a high-throughput high-content screen typically includes thousands to millions of genetic or chemical treatments. Each treatment is performed singly or repeated several times (biological or technical replicates), each replicate contains two or more fluorescent channels of microscopic images highlighting various biological concepts (cellular organelles or proteins), and each image comprises roughly 500-2000 pixels per dimension. The objective of analyzing this imaging data is to investigate how the treatments affect the cells and whether treatments cause similar or diverse effects. In computational terms, we aim to cluster the treatments based on the microscopic images, such that treatments that cause similar cellular changes group together.

As genetic or chemical treatments can have diverse effects and various cellular systems may respond differently, a high content screening data set contains many image classes which are not possible to define a priori. There is also unknown biological and technical noise in the data (e.g., individual cells not behaving consistently or issues in staining or image acquisition). These challenges plus the sheer volume and complexity of the imaging data makes data analysis and interpretation a daunting task, which has been an active research topic and has seen substantial development over the years^4–12^. Recent breakthroughs in deep neural networks have transformed data analysis^13^ and have been successfully pioneered in biomedical applications, particularly in the computer vision domain^14^. High-throughput high-content cellular imaging presents a unique opportunity and challenge for deep neural networks, with diverse innovative methods applied recently^15–22^. Although all these methods, deep neural networks based or not, are successful in analyzing their corresponding data sets, the strengths and limitations of each method are not clear since they have not been benchmarked in the same context. Thus, it remains an open question which method one should apply given a new high-content cellular imaging project.

In this study, we have implemented seven leading unsupervised image analysis methods for high-content cellular imaging assays. The cellular feature (CF) method segments cells and measures a suite of pre-defined features^11,23^. The image-level feature (IF) method measures a number of pre-defined features within the whole image without cell segmentation^24,25^. The pseudo-classification (PC) method defines individual treatments as pseudo-classes and feeds them into a neural network model for classification training to generate embeddings^17^. The metadata-guided learning (MGL) method relies on metadata to develop representation learning^22^. The deep clustering (DC) method iteratively uses embeddings to cluster and uses cluster labels to perform classification and generate embeddings^19^. The invariant information clustering (IIC) method trains a representation model by maximizing the mutual information between paired samples^26^. The transfer learning (TL) method applies a model pretrained from ImageNet directly on cellular images and extracts embeddings^16^. The above methods have been previously developed either for unsupervised learning generally or for high-content cellular images specifically, and all have showed promising results.

For the benchmarking data set we used the image set BBBC021v1^27^, publicly available from the Broad Bioimage Benchmark Collection^28^. In contrast to previous benchmarking efforts, manual annotation was not provided to the methods, but rather used for evaluation criteria afterwards. Each method showed strength and limitations in predicting compound mechanism of action (MoA), clustering similar compounds, and detecting novel phenotypes, and not one method outperformed the others across all metrics. We further investigated the predictions that disagreed with the manual annotations and the novel phenotypes detected as outlier by the methods, as well as examined the biological concepts of features automatically learned by deep neural networks

## Results

### Workflow and benchmark design

In order to have a fair and realistic evaluation, we applied our analysis methods on the BBBC021 benchmark dataset in an unsupervised manner and used identical evaluation criteria (**Fig. 1**). The BBBC021 dataset is the result of profiling 113 low molecular weight compounds across 8 concentrations in MCF-7 breast cancer cells labeled for DNA, F-actin, and β-tubulin^27,28^. A subset with 103 compound-concentrations from 38 compounds has been manually categorized into one of 12 MoAs based on visual inspection of the images or previous reports in the literature. This labeled subset served as ground truth for evaluation of the computational analysis predictions, similar to previous studies to train supervised or unsupervised models^11,16,18,22^. Since an actual screening campaign would not have the convenience of an annotated subset, we formulated the task in our benchmarking study such that each method was blinded to the MoA annotations as well as which images possessed them. Instead, the complete collection of images was made available to each method, along with the associated metadata such as compound name, concentration, plate and well identifiers (**Fig. 1**).

**Figure 1.**
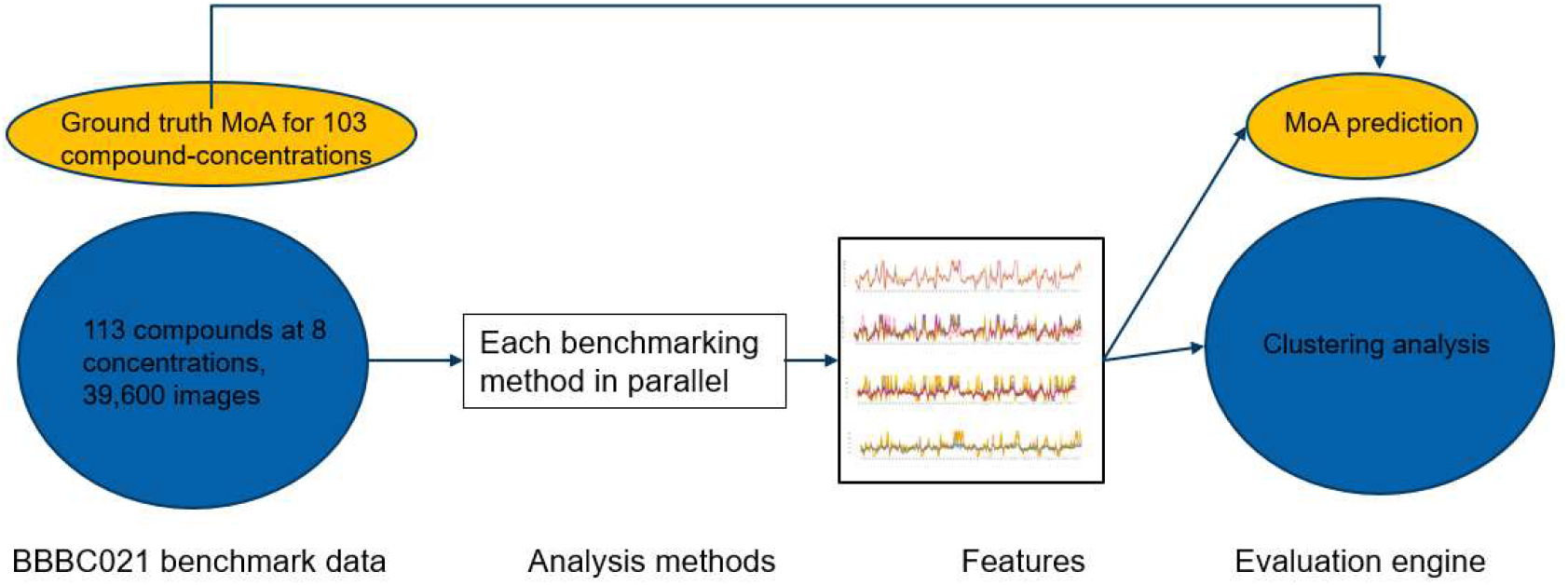
Schematic workflow of the benchmarking effort. The BBBC021 benchmark dataset has in total 113 compounds at 8 concentrations. A subset of 103 compound-concentrations (treatments) have manually annotated MoAs, which are used as ground truth. In parallel, each benchmarking method takes the total image dataset without the MoA annotations as input, and output features (embeddings for neural network-based methods and pre-defined features for feature engineering-based methods). The results are assessed by the evaluation engine using two criteria: MoA prediction accuracy for the ground truth annotated subset; clustering quality for the full dataset.

Each method and respective workflow generate a numeric feature vector (predefined features or neural network embeddings) for each image (see **Methods** and **Supplementary information** for details). We hypothesized that the feature vectors capture the cellular image phenotypes produced by the compound treatments and associated MoA, and evaluated their accuracy both in predicting MoAs and grouping compounds with similar MoAs.

### Classification results

We designed a simple 1-nearest neighbor (1-NN) classifier using cosine distance to evaluate the feature vectors generated by each method (see **Methods** for details). The NSC (not-the-same-compound) and NSB (not-the-same-compound and batch) accuracy values are shown in **Table 1**. The NSC accuracy ranges from 0.7 to 0.99 with a median of 0.94. The IF method has the lowest accuracy at 0.7 while the CF, DC, IIC and TL methods achieve >0.9 accuracy. The NSB accuracy is lower than NSC (range: 0.57-0.93, median: 0.79), which could be due to batch variation or a smaller sample size. The CF method suffers the highest decrease from the NSC accuracy to NSB. Overall, the DC, IIC and TL methods have the highest accuracy values across both metrics, showing their ability to capture cellular phenotypic information from the images.

**Table 1.** Summary of the evaluation results of classification and clustering tasks. NSC: not-same-compound accuracy; NSB: not-same-compound-and-batch accuracy; ARI: adjusted rand index; AMI: adjusted mutual information; sil_embedding: silhouette coefficient embedding; sil_clustering: silhouette coefficient clustering.

| Method | NSC<br>accuracy | NSB<br>accuracy | ARI | AMI | sil_embedding | sil_clustering |
| --- | --- | --- | --- | --- | --- | --- |
| Cell-level feature (CF) | 0.94 | 0.75 | 0.45 | 0.65 | 0.21 | -0.30 |
| Image-level feature (IF) | 0.70 | 0.57 | 0.39 | 0.52 | 0.10 | +0.06 |
| Pseudo-classification<br>(PC) | 0.84 | 0.73 | 0.54 | 0.64 | 0.21 | +0.11 |
| Metadata-Guided<br>Learning (MGL) | 0.89 | 0.83 | 0.51 | 0.69 | 0.30 | +0.02 |
| DeepCluster (DC) | 0.99 | 0.93 | 0.75 | 0.82 | 0.32 | +0.04 |
| Invariant information<br>clustering (IIC) | 0.96 | 0.90 | 0.81 | 0.81 | 0.42 | +0.36 |
| Transfer learning (TL) | 0.98 | 0.92 | 0.77 | 0.84 | 0.29 | +0.06 |

To further investigate how methods agree with each other, we list all NSC predictions from each method (**Fig. 2a**, agreement with manual annotation in green and disagreement in red) as well as the summary of all methods (**Fig. 2a**, grayscale based on the number of methods that disagree with the manual annotation). All methods are largely in agreement with each other and with the manual annotation, as shown by the green block. Interestingly, there are four treatments where four or more methods disagree with the manual annotation (predictions in **Fig. 2b**, example images in **Fig. 2c**). For colchicine at 0.03µM, the manual annotation is Microtubule destabilizers for which all methods disagree except for DC. The methods also disagree with each other, with four other different outcomes (Microtubule stabilizers, Kinase inhibitor, DNA damage, Cholesterol-lowering). This treatment was also reported previously where images in this class seem to have high levels of variation^17^. For cytochalasin D at 0.3µM, the three top-performing methods match the manual annotation (Actin disruptors) while the other methods disagree. However, for epothilone B at 0.1µM and latrunculin B at 1.0µM, a different set of methods agree with the manual annotation. Thus, there appears to be no single method that outperforms all others across all treatments, and an ensemble approach will not necessarily improve the overall accuracy.

**Figure 2.**
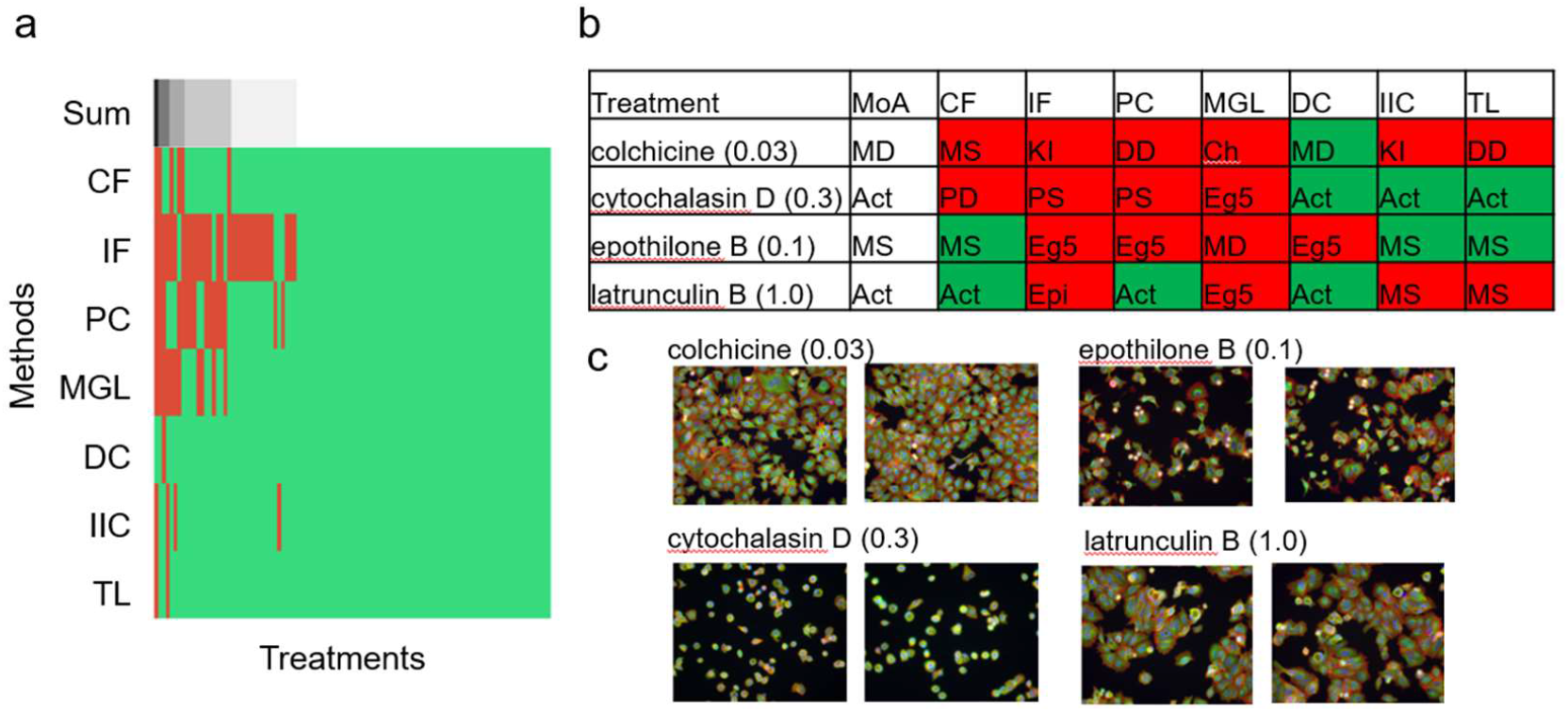
MoA prediction evaluation results. **a**, Each of the colored row represents the classification results of each method with 103 treatments on the horizontal axis. A correct prediction is labeled in green while a false prediction is labeled in red. The top black and white row is the sum of all methods. **b**, Top two treatments where most methods make false predictions, i.e., left most two lines from panel a. Color coding is the same as in panel a. **c**, Example images of the two treatments in panel b.

### Clustering results

For IIC and DC, the cluster labels were derived directly from the network output. For the other methods, standard clustering algorithms were run over the feature vectors to obtain cluster assignments (see **Methods**). We compared four metrics over the clustering results (**Table 1**). The adjusted rand index (ARI) and adjusted mutual information (AMI) evaluate the similarity between the predicted cluster assignments and the ground truth MoA. The ARI ranges from 0.45-0.81 while the AMI ranges from 0.52-0.84. The IIC, DC and TL methods are among the first performance tier with an average ARI and AMI around 0.8. The CF, IF, PC, and MGL are among the second tier with the average ARI and AMI as 0.3 and 0.2 lower than the first tier, respectively. The silhouette coefficient with predicted cluster labels (denoted by sil_coef_clustering) measures how compact the clusters are. The score ranges from 0.1 to 0.42, with IIC performing better than the others. The silhouette coefficient with ground truth MoA labels (denoted by sil_coef_embedding) evaluates how tight the embeddings are within each MoA class. Here, the coefficient had a larger range (−0.3 to +0.36) but IIC again outperformed the others.

### Novel phenotype detection

Each method attempted to detect novel phenotypes, defined as treatments different from any of the manually annotated MoAs. For this experiment, the similarity and clustering analysis were not standardized across methods; therefore, each method applied differing amounts of outlier stringency and the number of treatments labeled as novel varied for each method (see **Methods** and **Supplementary information** for details). However, multiple methods agreed on novelty for some treatments (**Fig. 3a**). Manual review indicated that these treatments were predominantly toxic phenotypes. For example, filipin at 1 – 10µM was labelled by most methods (six of seven) as novel; no cells were visible in the associated images, suggesting a strong toxic phenotype (**Fig. 3b**). Similarly, doxorubicin at 1 – 10µM, was labelled by four out of seven methods as novel. Here, the observation was a reduction in cell count along with changes in morphology, suggesting an unknown phenotype with slightly toxic effect.

**Figure 3.**
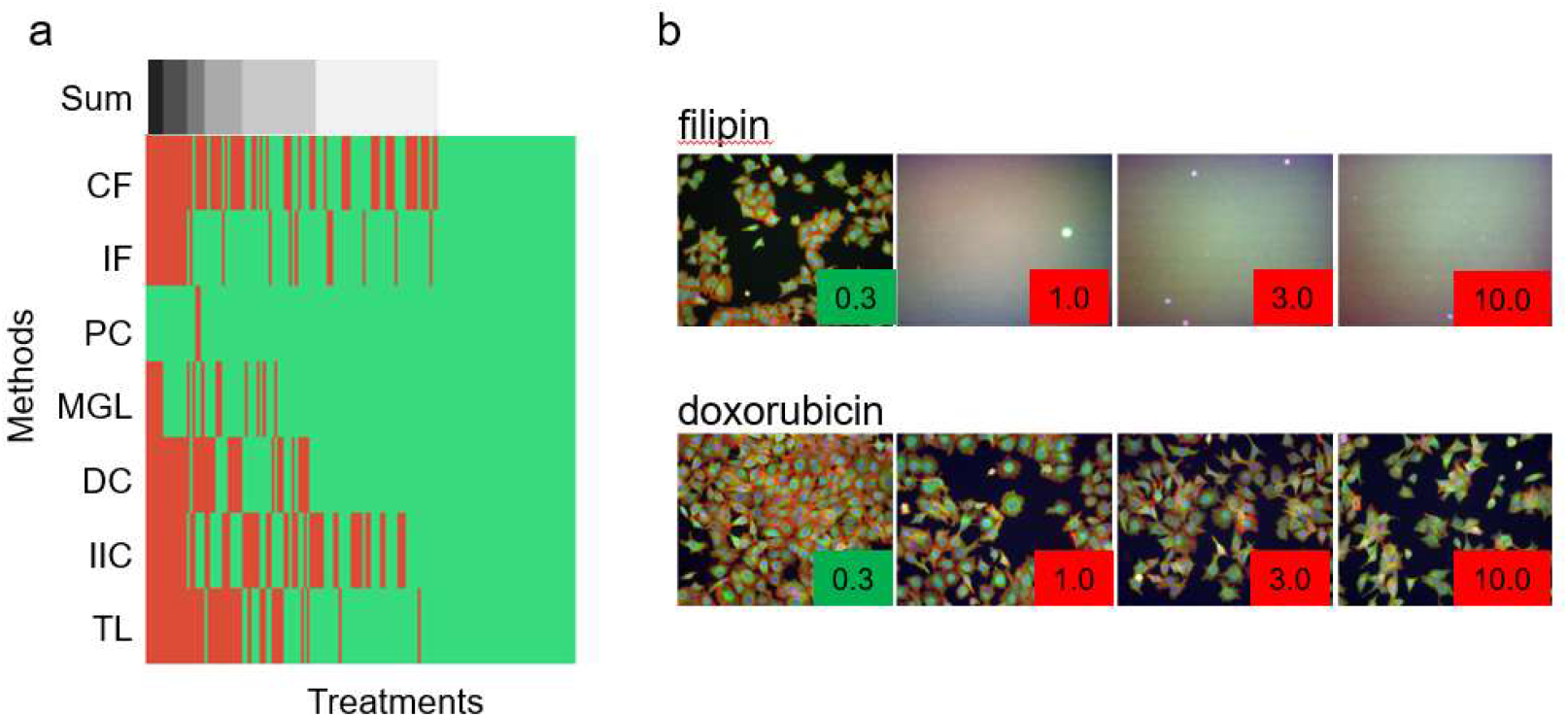
Novel phenotype detection analysis of BBBC021. **a**, Each row represents the outliers detected by each method, with methods on the vertical axis and the 906 treatments on the horizontal axis. “Novel” phenotypes are labeled as red, with remainder as green. The top grayscale bar illustrates the total number of methods labeling the same treatment found as “novel.” **b**, Examples of detected novel phenotypes.

### Feature interpretation

Contrary to CF and IF which use pre-defined features, methods based on deep neural networks learn the features automatically. To understand what features were learned, we used the raw features (without post-processing) from the CF method as a reference since they are widely used features defined by a priori biology knowledge. For every method, correlation similarity was calculated between each feature and each of the reference features. The distributions of the similarity values are shown in **Fig. 4a** as violin plots. As the reference feature set is a collection of cellular morphology, intensity and texture features across biological compartments and imaging channels, the similarity values of the reference set against itself have a large numeric range (0 to 1) with an average of 0.30. The post-processing steps of the CF method yields lower similarity values (range of 0 to 0.57 with an average of 0.14) as compared to the reference set. However, both the CF and IF methods have relatively higher similarity values with an average between 0.14 and 0.15 as compared with the deep neural network methods with an average between 0.10 and 0.13 which suggests that the neural network embeddings include human-interpretable features, at least in part. We investigated further by taking features from the TL method as an example and compared how compounds from the twelve manually annotated MoAs plus DMSO scored for each feature dimension. Some features clearly distinguish different MoAs (examples shown in **Fig. 4c**). We also plotted two features against each other for the whole data set (**Fig. 4b**) with example images for high and low values. Although the features could capture cellular intensity, object count or other texture measurements and it is impossible to pinpoint what exactly each feature represents, we can visually discern the high/low images, supporting that the learned features correlate with visual inspection and separate cellular phenotypes.

**Figure 4.**
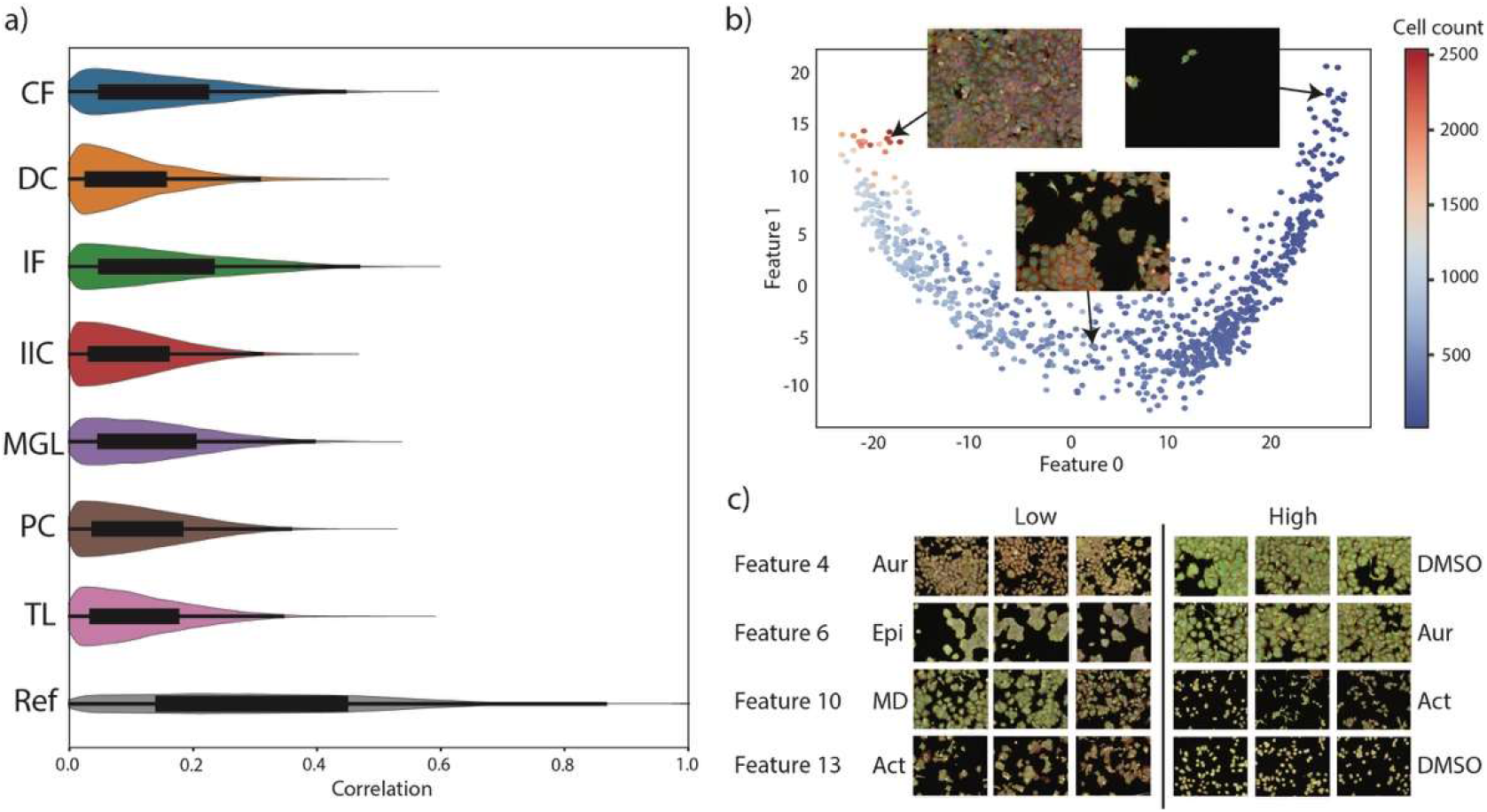
Interpretation of features learned by deep neural networks. **a**, Distribution of pair-wise correlation of features from each method against raw cellular features as the reference. **b**, 2D scatter plot of two features from the TL method and example images of high and low values. **c**, Selected features from the TL method and example images from MoA classes with high and low values of each feature.

## Discussion

Recent advancement of deep neural network methods has transformed how we analyze cellular images^29^, and has in principle solved tasks such as segmentation^30,31^ and modality translation^32,33^. However, the ultimate task, to understand the phenotypes captured by cellular images, remains elusive. Although various methods have been proposed, they were evaluated individually and often with unrealistic settings and criteria. Here we define the task as unsupervised representation learning to facilitate downstream classification and clustering of treatment mode of action. Importantly manual phenotype annotations are not provided to the methods, but rather used for evaluation criteria afterwards, which is similar to a realistic screening project. We benchmark five published deep neural network methods together with two classical feature-engineering methods, with consistent evaluation on a single well-characterized data set.

In the classification evaluation, DC performs the best with TL and IIC are closely following. The traditional CF method also predicts the MoA labels with a high accuracy but suffers more from batch effects. Interestingly, the methods disagree with the manual annotations consistently on some treatments suggesting potential issues in the imaging data or in the manual annotation. In the clustering evaluation, IIC, TL and DC perform the best on the ARI and AMI metrics while IIC leads on the silhouette metrics. All the methods can also detect novel phenotypes that are not in the manual annotated subset. The most novel phenotypes are images with artifacts or toxic treatments, which might be a limitation of this particular data set.

These methods vary substantially on computational resource and time needed. The PC, DC, IIC and MGL methods all need heavy computation resources proportional to the number of treatments and are thus not practical on larger data sets (∼100K treatments), while CF, IF and TL are scalable. All deep neural network-based methods require multiple graphics processing units to accelerate computation.

There are methods not included in this benchmark effort and new methods are being developed regularly. We have implemented methods we are familiar with and optimized them to the best of our knowledge. It is possible that certain methods can be further optimized or other methods can perform better. With the diversity and complexity of cellular imaging data, performance may differ depending on the data set and thus the presented benchmarking results only represent current implementations upon the BBBC021 data set. Further work is needed to benchmark promising methods on diverse data sets, especially as additional data sets have been published more recently^34,35^.

As deep neural network-based methods learn their embeddings automatically, it is important to understand the biological concepts behind these learned features, not only to investigate how the neural network works, but also to potentially bring novel biological insights. By performing correlation analysis between learned features and predefined features, our results suggest that the learned features contain not only information overlapping with predefined features but also new information. We hope this intriguing finding can trigger more research on this promising direction.

## Acknowledgements

We would like to thank Christopher Ball, Imtiaz Hossain, William Godinez and Florian Fuchs for their contributions.

## Methods

### Benchmark dataset

We used the image set BBBC021v1^27^ publicly available from the Broad Bioimage Benchmark Collection^28^. A detailed description of the dataset can be found on https://bbbc.broadinstitute.org/BBBC021.

### Image analysis methods

Seven methods were implemented for benchmarking: Cell-level feature analysis (CF); Image-level feature analysis (IF); DeepCluster (DC); Invariant information clustering (IIC); Metadata-Guided Learning (MGL); Pseudo-classification (PC); Transfer learning (TL). The specific details for each implementation are provided in the **Supplementary information**.

### Evaluation methods

To fairly benchmark the methods in this study, we blinded the methods to the MoA annotations in the BBBC021 dataset. Each method built models and feature embedding in an unsupervised manner based on remaining treatment metadata and submitted the embedding and cluster assignments per treatment for the full dataset. The predictions and associated compound and concentration metadata were then provided as input into a built-for-purpose evaluation engine. The engine ran two tasks for each input set and generated the following scores:

### Prediction of the MoA label for a given treatment

The evaluation was performed using the annotated ground truth subset of BBBC021. Given a treatment from a submission, the engine trained a 1-NN classifier with cosine distance over a subset of the feature vectors and the corresponding ground truth MoA. The subset was chosen based on two criteria: 1) NSC (not the same compound) which excludes the features from all concentrations of the same compound; 2) NSB (not the same compound and batch) which further excludes any compounds from the same batch. For NSB, two MoAs (Cholesterol lowering and Kinase inhibitors) were removed as they only appear in a single batch. The trained classifiers were then applied to the query treatment to obtain the NSC and NSB predictions. This process was iteratively applied to all treatments, which were then finally compared to the ground truth to obtain the NSC-accuracy and NSB-accuracy.

### Clustering of compounds based on MoA

Four benchmarking scores were used for evaluation of predicted cluster quality:

#### Adjusted rand index (ARI) and adjusted mutual information (AMI)

These metrics were used to evaluate the similarity between the predicted cluster structure and ground truth, ignoring permutations. Since the two scores require knowledge of the ground truth classes, they can only be applied to the annotated subset. More specifically, the rand index is the percentage of sample pairs assigned to the same cluster in both the predicted and ground truth clusters. The ARI is then the corrected-for-chance version of rand index, defined as 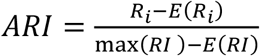. Formally, if there are *n* elements with two partitions, e.g., {*X*_1_, …, *X*_*r*_} and {*Y*_1_, …, *Y*_*s*_}, the ARI is calculated as

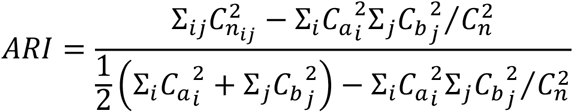

where *n*_*ij*_ = |*X*_*i*_ ∩ *Y*_*j*_ |, *a*_*i*_ = ∑_*j*_ *n*_*ij*_, *b*_*j*_ = ∑_*i*_ *n*_*ij*_.

The adjusted version AMI is calculated as

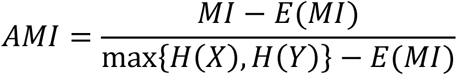

where the mutual information *MI* between two partitions is defined as

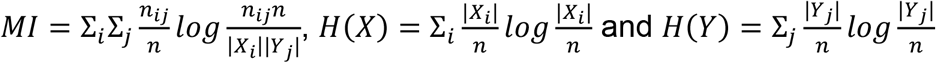

#### Silhouette coefficient of predicted feature clusters versus predicted MoA and ground truth MoA clusters

The silhouette coefficient measures the compactness of a cluster as compared to neighboring clusters, and is defined as is (*b* − *a*)/max (*a, b*) where *a* is the mean intra-cluster distance and *b* is the mean nearest-cluster distance. The silhouette coefficient was computed to compare the predicted feature vectors with respect to both the predicted MoA clustering and the ground truth MoA clustering. The former provides a measure of how separable the MoA clusters predicted by the embeddings were, whereas the latter measures how well the ground truth MoA clustered in the embedding space.

