## Supplementary information for "High-content cellular screen image analysis benchmark study"

[1]Authors collaborated on the project as a team; names are ordered alphabetically.

[2]Novartis Institutes for BioMedical Research Inc., Cambridge, MA, USA

[3]Novartis Institutes for BioMedical Research, Basel, Switzerland

[4]Present address: Children's Cancer Institute, Lowy Cancer Research Centre and School of Women and Children's Health, Faculty of Medicine and Health, UNSW Sydney, NSW, 2031, Australia

### Supplementary information

### Supplementary methods

#### Cell-level feature analysis (CF)

**Summary** In contrast to plate reader technologies that produce a single readout per well, single-cell analysis allows for a rich set of multiplexed readouts collected from individual cells and their associated sub-compartments. The resultant profile is well suited for connecting subtle morphological phenotypes back to the originating image feature and cellular compartment that produced the phenotypic change. The key elements of the analysis workflow have been described extensively elsewhere, for example in[1].

**Specifics**

1. Non-uniform illumination can introduce distortions into the raw fluorescence images, biasing the resultant profiles. A retrospective illumination approach is used to correct the images on a per-channel, per-plate basis using the CellProfiler open-source image-analysis package[2,3].
2. The nuclei, associated cell body and cytoplasm are segmented from the corrected images via the standard model-based approach in CellProfiler, which requires optimization by visual inspection.
3. From each nucleus, cell and cytoplasm object, a variety of image features are extracted including shape, intensity, texture, cross-channel correlation and spatial context metrics. The embedding formed from these features describes the phenotypic state of each cell.
4. The element-wise median of the replicate profiles is computed to generate the final embedding for each treatment[4]. The resultant feature vector is then further processed using the following steps:
    - Removal of spatial plate-position artifacts using local linear regression at the per-feature and per-plate level.
    - Each feature is robust Z-scored (e.g., median/MAD instead of mean/standard deviation) using the feature distributions of the DMSO wells.
    - Since not all features contain useful information, the features are filtered to remove those with low variance (defined as rejecting features where the number of unique (distinct) values divided by number of wells < 10%),
    - Features which are highly correlated and/or nearly collinear are also rejected to yield the final feature vector.

#### Image-level feature analysis (IF)

**Summary** Although segmentation-based image analysis has dominated the high content screening community, morphological features can be successfully extracted and measured from the whole image without segmentation. The image-level classification methodology used in this study is based on previous work by the Broad Institute Imaging Platform[5]. It uses a set of published image analysis modules (collectively referred to as CP-CHARM) for the popular CellProfiler open-source software[6]. The CP-CHARM pipeline extracts a wide variety of morphological image features without cell/object segmentation. The extracted multi-scale features (typically after dimensionality reduction) are used to classify the images. Major advantages of this methodology include minimal need for pipeline and parameter optimization, relatively well-understood morphological features and ease of implementation through CellProfiler. The CP-CHARM image classification methodology used in this study is adapted from previous work described in the development of the Fluopack screening platform[7].

**Specifics**

1. CellProfiler feature extraction. Each BBBC021 image is acquired at 3 different channels. Images from each channel are processed with the CP-CHARM pipeline to extract multi-scale morphological features (high-contrast features, polynomial decompositions, pixel statistics, textures). The combined 3 channel feature vector is comprised of 2904 features.
2. Feature reduction and annotation. CHARM features are analyzed using the NIBR Multi-parametric Data Analysis (MPDA) system that performs feature analyses for correlation, collinearity and variance. MPDA selects features that are deemed most informative for discriminating positive and negative control images. Single value features or features with low information, large correlation and high collinearity are excluded. The resulting three-channel feature vector is composed of 686 features. These features are exported as a CSV file and BBBC021 treatment metadata columns are added using simple SQL manipulations.
3. Batch normalization. A custom R script is used to normalize the CHARM features per BBBC021 batch. For this, we use the classification and regression training *caret* R package and specifically the *preProcess* S3 method with sequential "center", "scale" operations.
4. Cluster assignment. We used K-means clustering in R to assign treatments (compound/dose combinations) to clusters using the batch normalized features. We estimated the right number of clusters using the *FitKMeans* function in the R *useful* package. The plot of the ‘Hardigan’s rule’ suggested 18 centers for the K-Means algorithm. We obtained the final assignment of treatment to clusters running K-means with *centers=18* and *nstart=25*.
5. Novelty Detection. To enable novelty detection, after cluster assignment, we added to each treatment tuple an additional column GT that was assigned values as follows: ‘GT’ if the treatment belonged to the ground truth set; ‘KNOWN’ if the

compound MOA is known; ‘UNKNOWN’ if the compound MOA is unknown. We then selected clusters for treatment tuples that did not include any ‘GT’ treatments. Four clusters with 127,4,8,5 treatments (compound/dose combinations) were identified. The treatments in each of these clusters were considered novel since they did not include any of the 103 ground truth treatments.

#### Pseudo-classification (PC)

**Summary** The pseudo-classification method uses technical labels instead of manual ground truth labels, and thus converts the unsupervised task into a supervised task. The embeddings are subsequently extracted and used for downstream analysis. The method is first published in[8] and adopted for cellular images in[9].

**Specifics**

1. Preprocessing. For each channel and each plate, the control (DMSO & taxol) columns were selected for intensity normalization. The 0.25 and 99.75 percentile intensity was calculated, and used to linearly scale the remaining images from that computed range to [0.0-1.0]. Values outsize this range were clipped.
2. Training. The normalized non-control images were assigned to a pseudo-class based on the treatment and concentration used, and these labels were used to train a softmax classifier using cross-entropy loss. All the setup was the same as in[9].
3. Clustering. The penultimate layer outputs from the trained model were used without post-processing for cluster analysis with K-means, using k=100.

#### Metadata-Guided Learning (MGL)

**Summary** We develop a novel approach to visual representation learning that is guided by metadata (see previous publication[10]). In high-context-screening, metadata can typically be derived from the experimental layout, which links each cell image of a particular assay to the tested chemical compound and corresponding compound concentration. In general, there exists a one-to-many relationship between phenotype and compound since various molecules and different dosage can lead to one and the same alterations in biological cells.

**Specifics**

1. Preprocessing. As an input we assume BBBC021 cell images with their corresponding metadata, giving information about the applied compound and tested concentration. The images were illumination corrected to address differences in light conditions during measurement. Due to memory constraints, the images were furthermore resized by factor two, so that our algorithm can

process more images in one batch. Moreover, all image values were normalized, making it easier for the machine to learn from the data.

2. Sampling. Our idea of using metadata to guide our learning algorithm is reflected in the way we construct batches. In our approach a batch contains multiple images tuples, where each tuple consists of two images with the same metadata or rather treatment, referring to the same compound and concentration. Furthermore, all the tuples in one batch have metadata that is different from each other. In that sense, the metadata of a typical batch with 8 tuples would look like the following:

| **Tuple** | **1** | **2** | **3** | **...** | **8** |
|---|---|---|---|---|---|
| **Compound** | Acyclovir | Bleomycin | Acyclovir | ... | Taxol |
| **Concentration** | 0.1 µmol | 1.5 µmol | 0.5 µmol | ... | 0.3 µmol |

It is worthwhile mentioning that we can oversample, since our dataset contains 906 distinct treatments, leading to countless possible combinations of image tuples with different metadata. More precisely, we sample 4000 batches, each with 8 tuples and 16 images respectively. Doing the math, we generate 64'000 data points out of originally 13'200 observations, oversampling by factor five.

3. Loss function. Given a batch of image tuples with distinct metadata, we can now define a loss function that constraint our learning algorithm to extract image features that are similar for observations from the same tuple and dissimilar for images from different tuples. These constraints enforce a features space that guarantees spatial closeness for images with the same metadata and spatial remoteness for images with different metadata.

   Since we can create batches with tuples that have different metadata and still belong to the same phenotype, our learning algorithms occasionally ends up in a contradictory position, where it simultaneously tries to pull together and push away same class images. However, it is exactly these contradictions that lead to the reshaping of our feature space in a way that same class images with different metadata are clustered together after all.

4. Model architecture. We employed a ResNet50v2 architecture that was pre-trained on the ImageNet dataset and performed transfer learning to adjust the model parameters to our own domain. The original architecture was modified by removing the top layer, performing average pooling, and adding a l2-normalisation layer as well as a lower-dimensional projecting head. The projection head is only used during training and is cut off when making predictions. For back-propagation we used the Adam algorithm with standard parameters. Furthermore, we plugged in our described loss function, which computes the gradient for updating the model parameters.

   The selected model architecture was trained over 15 epochs. In each epoch the model processed a total amount 4000 batches, each containing 8 tuples or rather

16 images. Since we used 4 GPUs in parallel, our model was able to handle 4 batches with 64 images simultaneously, resulting in 1000 update steps per epoch. To keep all the 4 GPUs busy constantly, we employed 2 CPUs for preprocessing the images and passing them to the model on the fly.

5. Evaluation. After training the ResNet50v2 architecture, we remove the previously added projection head. The remaining convolutional neural network is used to embed our original image data into the learned feature space. The model takes our preprocessed images as input and outputs the corresponding feature vectors, which are 2048-dimensional and l2-normalized.

   Since our feature space is high-dimensional, we employ Principal Component Analysis (PCA) for dimensionality reduction. In order to remove noise, the resulting 128-dimensional feature vectors are furthermore corrected by transforming them into the feature space spanned by the DMSO control group.

   Given the reduced and corrected feature matrix for all images (13'200 x 128), we can now aggregate the information by computing the mean feature values for each of the 906 distinct treatments. The resulting matrix (960 x 128) exhibits one row for each compound and concentration combination and one column for each latent factor spanning the reduced and corrected feature space.

   As a final post-processing step, we group our treatment-level information by spectral clustering. The number of groups should equal the number of clusters or rather phenotypes presented in the original dataset. Now we are in a position to measure the performance of our approach by comparing our computed groups with the original clusters.

6. Hyper-Parameters. There exist a couple of hyper-parameters that influence our model performance. Although we did not evaluate their impact systematically, we still want to mention them at this point. A more thorough evaluation is planned in the future.

   In the image preprocessing step, the choice of resizing factor, normalization technique, and illumination correction makes a difference. In the sampling step, the number and size of batches play a critical role in how well our model learns and how fast the loss converges.

   Regarding our model architecture, the pooling type, normalization technique, and size of projection head tend to influence the results. Furthermore, the choice of optimization algorithm and its corresponding parameters makes a difference. Moreover, the number of training epochs has a direct impact on the performance. And of course, we could employ another architecture and loss function all together.

   In the post-processing step, the dimensionality reduction technique, the size of the reduced feature vectors, and the way of aggregating at treatment-level have an influence on the final clustering results. In addition, the choice of clustering algorithm has an impact on the overall model performance.

#### Deep Clustering (DC)

**Summary** The deep clustering framework is a self-supervised method previously published in[11] and adopted for cellular images[12]. For this benchmark study, we started from the UMM Discovery[12] and further optimized the method to get even better results.

**Specifics**

1. Model. We are using the on ImageNet[13] pretrained ResNet-50[14] architecture without the classification layer.
2. Preprocessing. All images were illumination corrected as in[15].
3. Model Training. One crop of each image at different scales is fed through the neural network to extract the corresponding features. After the feature embedding extraction of all the images, we do a well-level aggregation by the median. The idea behind this is that we assume that images from the same well, should look in average the same. Additionally, this speeds up the later t-SNE computation immensely. The well-level embeddings are batch corrected with Combat and TVN and preprocessed by t-SNE and L2 normalization. Next, the normalized embeddings are clustered with k-means. The cluster assignments from k-means are the new pseudo-labels and the number of clusters will become the size of the classification layer. After creation of the pseudo-labels, the training process of the network starts, where images are forwarded through the NN and the last classification layer to get a prediction for each image. The weights of the neural network are optimized by taking the cross-entropy loss function between the prediction and the pseudo-label and back-propagating it through neural network. This series of steps is repeated for each epoch.
4. Augmentation. We use on-the-fly augmentation of the input images during the training phase. The augmentations include resized crops, flipping, rotations. Additionally, we use CutMix[16] regularization within each batch during training. This regularization method should reduce the batch effect even further.
5. Model inference. After training, all preprocessed images are fed into the finetuned Resnet-50 without classification layer to generate the feature embeddings. As in 12, we use the combination of Combat with Typical variation normalization (TVN) to remove the batch effects.

#### Invariant information clustering analysis (IIC)

**Summary** Invariant information clustering (IIC) is an end-to-end image clustering algorithm proposed in[17], which has been validated to achieve state-of-the-art performance in sematic clustering of natural images. Here, we introduce how we adapt IIC to cellular images in the context of high content screening.

**Specifics**

1. Model**.** The IIC took image pairs as input, each pair contains images from the same treatment group. Then the image pairs were fed into a randomly initialized CNN classifier to generate cluster assignment probability of each image. We used ResNet34 to balance between performance and computational cost and set the number of clusters to be 50. The CNN was optimized by maximizing the mutual information of image pairs $I(\phi(x), \phi(x'))$. Given a batch of $n$ image pairs and number of clusters C, suppose $P = \frac{1}{n}\Sigma_i \phi(x_i)\phi(x_i')$, the mutual information loss can be calculated as
$$L_{MI} = -\Sigma_c \Sigma_{c'} P_{cc'} log \frac{P_{cc'}}{P_c P_{c'}}$$
where $P_c = \sum_{c'} P_{cc'}$ . In BBBC021 dataset, we have control wells (treated by DMSO) for all plates. Therefore, there are much more samples from the DMSO treatment, indicating much higher variance than other treatments. To deal with the imbalance, we constrained all DMSO images to be in the first 8 clusters (the order does not matter), by introducing an additional loss term:
$$L_{DMSO} = \Sigma_{x\ in\ DMSO} - \log(\max_{i \le 5} \phi(x)_i)$$
We then have the loss function for CNN training as
$$L = L_{MI} + \alpha L_{DMSO}$$

2. Augmentation**.** Within each batch, on-the-fly augmentation was applied, including random rotation and perturbations of normalization. More specifically, denote $Q(p)$ be the $p$th quantile of the pixel intensity of an image, we estimate
$$\mu = \frac{Q(25) + Q(75) + Q(50) * 2}{4}$$
$$\sigma = Q(75) - Q(25)$$
Then the image was firstly rotated by a random angle, and then centered by removing $r_1\mu$ and normalized by dividing $r_2\sigma$, where $r_1$ and $r_2$ are sample from $U[0.7,1,3]$. The image was finally resized to 280x224.

3. Model Training. To train the network, ADAM optimizer was employed with a learning rate of $10^{-4}$. Within each batch, we firstly use the standard distributed sampler to get 96 images from 8 GPUs. Then for each image, we randomly draw 5 images from the same treatment to create image pair containing the same original image. The sample repeat together with distributed learning (larger batch size) would stabilize the estimate of $L_{MI}$ within each batch. The training consists of 2400 epochs with randomly initialized Resnet34.
4. Model inference. After training, all the original images were fed into the Resnet34. The last layer and second to last layer were used as cluster labels and features, respectively. Novel clusters were defined by subsets of clusters of IIC (among a total of 50) to have no treatments with known MOAs.

#### Transfer learning (TL)

**Summary** The transfer learning approach uses state-of-the-art neural network architectures pretrained on big dataset consisting of consumer images to extract embeddings from new images.

##### Specifics

1. Model. In our case we use the on ImageNet[13] pretrained DenseNet121[18]. The classification layer of the neural network is removed to get feature embeddings for each image.
2. Preprocessing. All the images were illumination corrected as in[15].
3. Model Training. The pretrained model was directly used without further finetuning.
4. Model Inference. Each single-channel image is fed to the neural network separately. To fulfil the input dimensions (-1,3,224,224) of the neural network pre-trained on consumer images, copies of the same single-channel image are stacked on top of each other. Therefore, we extract for each single-channel image a feature embedding. The three embeddings for each of the single-channel images are concatenated together to get one long embedding for each field-of-view. Next, we perform a PCA dimensionality reduction to 128 dimensions followed by batch correction. We found that the combination of Combat with typical variation normalization (TVN) gave the best results as described in[12].
